# Resolvin D1 and D2 reduce SARS-Cov-2-induced inflammation in cystic fibrosis macrophages

**DOI:** 10.1101/2020.08.28.255463

**Authors:** Antonio Recchiuti, Sara Patruno, Domenico Mattoscio, Elisa Isopi, Antonella Pomilio, Alessia Lamolinara, Manuela Iezzi, Romina Pecce, Mario Romano

## Abstract

Resolvins (Rv) are endogenous lipid autacoids that mediate resolution of inflammation and bacterial infections. Their roles in SARS-CoV-2 and COVID-19 are of considerable interest in the context of cystic fibrosis (CF) given the paucity of data regarding the effect of this virus on immune cells from individuals with CF. Here, we provide evidence for Rv biosynthesis and regulatory actions on CF macrophage inflammatory responses.

## Introduction

The coronavirus disease 2019 (COVID-19) pandemic caused by the Severe Acute Respiratory Syndrome Coronavirus 2 (SARS-CoV-2) constitutes an unprecedented global health threat, with ∼ 22 mln confirmed cases and > 700,000 deaths worldwide. It is clear now that SARS-CoV-2 infections present with variable symptoms, with many people developing a severe pneumonia that progress towards respiratory distress syndrome, sepsis, respiratory failure, and multi-organ system dysfunction, whereas others show mild cough, fever or flu-like illness that resolve in a few days. Of interest, evidence signifies that the viral load is not correlated with the worsening of the symptoms, while cytokine storm, increase in inflammatory mediators and an imbalance in innate and adaptive immunity are associated with a poor prognosis ^1^. Risk factors for severe illness include comorbidities associated with unresolved inflammation, such as diabetes, cardiovascular diseases, cancer, chronic pulmonary diseases^2^, signifying that a defective resolution of inflammation can play important roles in the pathophysiology of COVID-19.

Cystic fibrosis (CF) is a multiorgan condition that includes chronic lung disease as well as systemic inflammation with high serum concentrations of inflammatory markers such interleukins (IL)-6, C-reactive protein, and ferritin^3,4^. Despite enormous strides have been made in the management of CF with the introduction of highly effective modulator therapies, unresolved inflammation and chronic infections remain constitutive in people with CF, as demonstrated by several longitudinal studies with patients taking CFTR modulators^5–7^. Since the early phases of the SARS-CoV-2 outbreak, people with CF have been considered at high risk for COVID-19 (13) because they have multi-organ chronic inflammation, bear persistent bacterial lung infections that may be exacerbated by a further viral stimulus, and have immune system dysfunctions involved in the response to CoV-2 and defective pro-resolution mechanisms that can determine an overshooting inflammatory reaction with detrimental consequences for health and life (18–21).

Resolution of inflammation is an active process introduced by the biosynthesis of specialized pro-resolving mediators (SPM), e.g., lipoxins (LX), resolvins (Rv), Protectins and maresins ^12^. SPM are evolutionarily conserved potent chemical signals that prevent excessive leukocyte infiltration and activation, balance inflammatory cytokines and chemokines, counter cytokine storm, regulate macrophage (MΦ) phenotype skewing, and protect inflamed organs from damage. SPM also act on lymphocyte maturation, T cell differentiation, and IgG switch (1) and enhance antimicrobial responses promoting resolution of infections, including bacterial and viral pneumonia (4–8). Hence, their actions on SARS-Cov-2 infection in CF are of interest

## Materials and Methods

### Chemicals

SARS-CoV-2 S1, S2, and N recombinant proteins were purchased from RayBiotech (Peachtree Corners, GA). RvD1, RvD2, LXA^4^, and RvD1 EIA were from Cayman Chemicals. RvD1 and RvD2 were stored and prepared before each experiment as previously published (6). Gibco cell culture media, fetal bovine serum (FBS), and supplements were purchased from Thermo Fisher Scientific. IL-8 ELISA kits were from Peprotech (London)

### MΦ culture and phagocytosis

Peripheral blood was obtained from volunteers with CF (age≥18 yrs; FEV1%≥70) that did not have exacerbations in the 4 weeks prior to blood collection (Prot. 2301/2017). Monocytes were grown onto plastic surfaces (Eppendorf, Milan) 7-10 days in 10 % FBS RPMI medium plus GM-CSF (10 ng/mL). A minimum of 24 h washout period was carried out to remove residual GM-CSF effects and cells were maintained in FBS-free or 1% FBS medium throughout the experiments.

### Luminex, RNA, and miRNA analyses

A Milliplex Magnetic Bead array kit (Merck Millipore, Milan) was used for measuring 30 cytokines, chemokines, interferons, and growth factors in cell-free media from experiments with MΦ. Total and small RNA (containing microRNA) were extracted from MΦ using the Roche High Pure miRNA Isolation kit (Roche, Milan) or the Quick-RNA MicroPrep from Zymo Research (Irvine, CA) and quantified using a UV nanophotometer. cDNA was reverse transcribed from 100-150 ng of total RNA using the SuperScript VILO Master Mix (Thermo Fisher Scientific) with ezDNAse treatment and used (1-5 ng/reaction) to assess gene expression with real time PCR as in ref. (22, 23). microRNAs were determined as previously published from 100 pg of cDNA synthesized with the miScript RT kit (Qiagen, Milan). Real time PCR were analyzed using the relative quantification method previously described (23).

### Stastistics

Results are reported as mean ± SE. Multiple comparisons were carried out with One Way ANOVA and Holm-Sidak or Dunn’s post-hoc test depending on variances among groups. P < 0.05 were considered statistically significant.

## Results

### Characterization of SARS-CoV-2 triggered responses by CF MΦ

The SARS-CoV-2 is an enveloped virus whose virion is composed of a phospholipid bilayer, covered by spike (S) proteins, which encloses the nucleocapsid made of a single stranded RNA and phosphorylated nucleocapsid (N) proteins. The S protein has the function of conveying binding of SARS-Cov-2 to target cells trough the binding to ACE2 receptors on host cell surfaces. Once bound to ACE2, the S protein is cleaved by the transmembrane Ser-protease 2 (TMPRSS2), which is essential for its priming, into the S1 and S2 subunits resulting in viral-host membrane fusion and virus endocytosis (24). On the contrary, the N protein binds to the virus RNA to ensure maintenance of a “beads-on-a-string” conformation and is, hence, essential for viral replication (25). To characterize host responses to CoV-2 in CF, monocyte-derived MΦ, which are key immune cells and contribute to the exaggerated non resolving inflammatory response in CF (26), were treated with CoV-2 S1, S2, or N proteins as surrogate of viral infection, and the release of cytokines, chemokines, IFN, and growth factors was determined. S1, S2, and N treatment resulted in a significant increase in IL-8 release by MΦ. Other chemokines involved in leukocyte recruitment were also significantly enhanced by CoV-2 although at a much lower extent: monocyte chemoattractant protein-(MCP)-1, macrophage inflammatory protein (MIP)-1α and 1β, and RANTES. On the contrary, IL-6, tumor necrosis factor (TNF)-α, IL1-β, and IFN-α and γ were not modified by CoV-2 and IL-1 receptor antagonist (IL1RA) was the only protein reduced by S1, S2, and N proteins in MΦ (Fig. 1A and Table 1). Gene expression analyses revealed that mRNA levels of IL-8 and IFN-α, β, and γ were not modified by S1, whereas MIP-1α, TNF-α, and IL-6 transcripts were significantly and strongly (∼ 100 -300 fold *vs* baseline in untreated cells) upregulated and RANTES was increased, although it did not reach significance (Fig. 1B). Collectively, these results indicate that CoV-2 S and N proteins have a direct, infection-independent pro-inflammatory action of CF MΦ, with protein N being a more potent inflammation stimulus.

**Table 1.** Mediators of innate and adaptive immunity released by CF MΦ in response to CoV-2.

|  | Ctrl |  | S1 |  |  | S2 |  |  | N |  |  |
| --- | --- | --- | --- | --- | --- | --- | --- | --- | --- | --- | --- |
|  | Mean | SE | Mean | SE | P vs Ctrl | Mean | SE | P vs Ctrl | Mean | SE | P vs Ctrl |
| IL-8 | 2.1 | 0.7 | 1516.0 | 919.4 | * | 2537.6 | 2160.9 | * | 9068.0 | 7822.8 | * |
| IL-1RA | 3092.8 | 353.4 | 2234.3 | 52.7 | * | 2230.3 | 274.5 | ns | 1994.5 | 576.9 | ns |
| GM-CSF | 0.0 | 0.0 | 6.2 | 3.2 | * | 10.7 | 6.2 | * | 3.7 | 0.7 | * |
| MCP-1 | 10.2 | 4.5 | 107.6 | 24.5 | * | 39.4 | 23.8 | * | 48.9 | 26.1 | ns |
| MIP-1 alpha | 14.6 | 10.4 | 532.0 | 58.8 | * | 283.2 | 89.4 | * | 356.9 | 201.1 | * |
| MIP-1 beta | 4.4 | 4.1 | 292.4 | 21.6 | *** | 148.4 | 11.1 | 0.001 | 373.4 | 52.7 | * |
| RANTES | 10.1 | 0.5 | 16.7 | 5.0 | * | 16.5 | 4.5 | * | 18.4 | 4.9 | * |
| TNF alpha | nd | - | nd | - | - | nd | - | - | nd | - | - |
| VEGF | nd | - | nd | - | - | nd | - | - | nd | - | - |
| EGF | nd | - | nd | - | - | nd | - | - | nd | - | - |
| EOTAXIN | nd | - | nd | - | - | nd | - | - | nd | - | - |
| FGF-2 | nd | - | nd | - | - | nd | - | - | nd | - | - |
| G-CSF | nd | - | nd | - | - | nd | - | - | nd | - | - |
| HGF | nd | - | nd | - | - | nd | - | - | nd | - | - |
| IFN alpha | nd | - | nd | - | - | nd | - | - | nd | - | - |
| IFN gamma | nd | - | nd | - | - | nd | - | - | nd | - | - |
| IL-1 beta | nd | - | nd | - | - | nd | - | - | nd | - | - |
| IL-10 | nd | - | nd | - | - | nd | - | - | nd | - | - |
| IL-12 | nd | - | nd | - | - | nd | - | - | nd | - | - |
| IL-13 | nd | - | nd | - | - | nd | - | - | nd | - | - |
| IL-15 | nd | - | nd | - | - | nd | - | - | nd | - | - |
| IL-17A | nd | - | nd | - | - | nd | - | - | nd | - | - |
| IL-2 | nd | - | nd | - | - | nd | - | - | nd | - | - |
| IL-2R | nd | - | nd | - | - | nd | - | - | nd | - | - |
| IL-4 | nd | - | nd | - | - | nd | - | - | nd | - | - |
| IL-5 | nd | - | nd | - | - | nd | - | - | nd | - | - |
| IL-6 | nd | - | nd | - | - | nd | - | - | nd | - | - |
| IL-7 | nd | - | nd | - | - | nd | - | - | nd | - | - |
| IP-10 | nd | - | nd | - | - | nd | - | - | nd | - | - |
| MIG | nd | - | nd | - | - | nd | - | - | nd | - | - |

Concentrations of proteins involved in innate and adaptive immune response released by monocyte-derived MΦ isolated from volunteers with CF treated (3 h, 37 °C) with recombinant CoV-2 spike proteins subunit 1 (S1), subunit 2 (S2), or nucleocapsid protein (N) at 10 µg/mL. Protein concentrations were measured using a multiplex Luminex array kit. Results are reported as pg/mL and are of experiments with cells from 4 different donors. nd, not detected (below limit). \*, P < 0.05; \*\*, P < 0.01.

**Figure 1.**
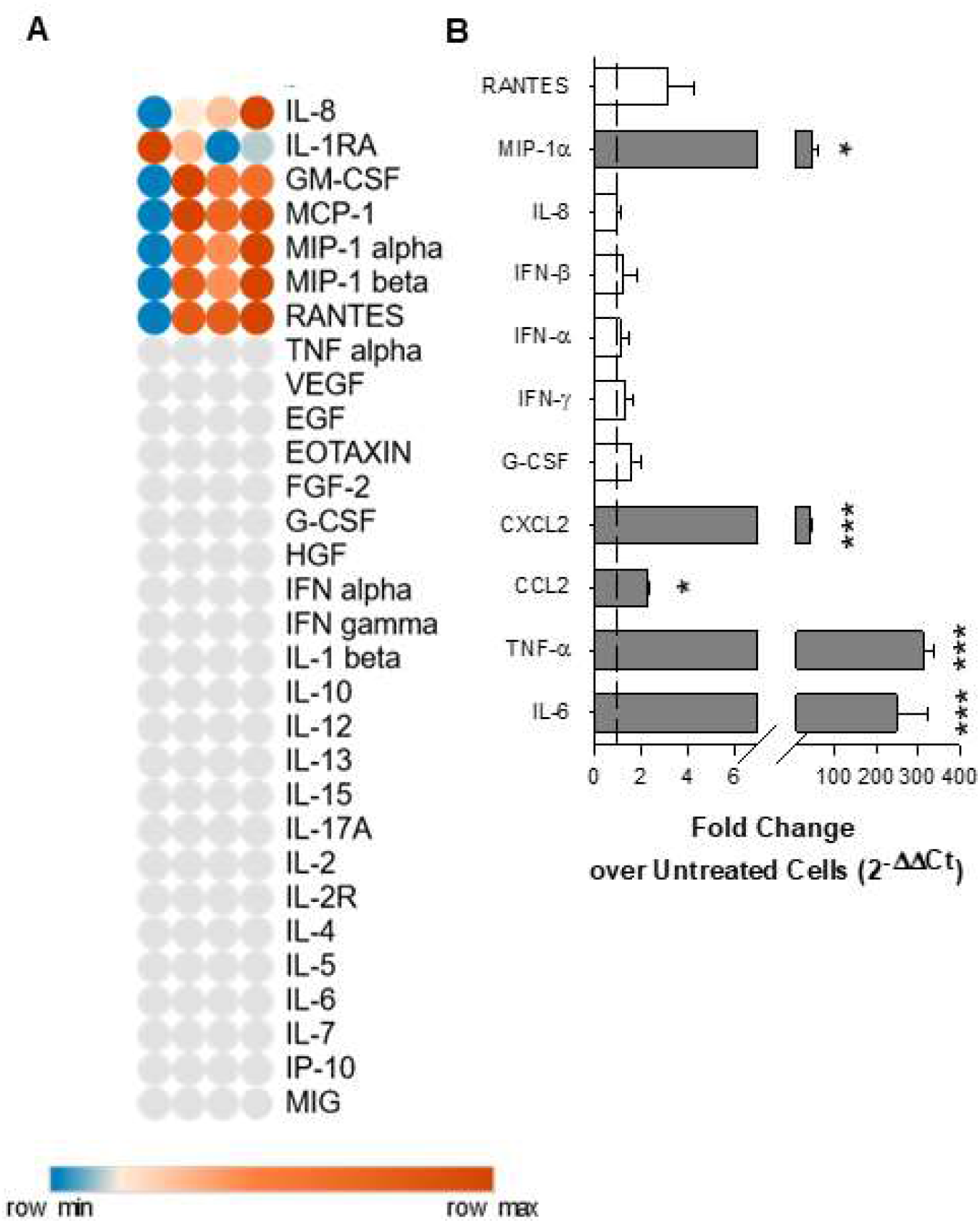
Inflammatory mediators regulated in CF MΦ in response to SARS-CoV-2 proteins. Monocyte-derived MΦ collected from volunteers with CF were stimulated with S1, S2, or N proteins (10 µg/mL, 3h). Protein in supernatants **(A)** and mRNA **(B)** levels of genes involved in innate and adaptive immunity were measured using a multiplex Luminex array kit (Millipore) or real time PCR. Results are mean ± SE of experiments with cells from 4 different donors. *, P < 0.05; **, P < 0.01; ***, P < 0.001 vs untreated cells.

### SARS-CoV-2 activates RvD1 biosynthesis

SPM are biosynthesized in inflammatory reactions following infection (6, 8, 23). In order to determine if SARS-CoV-2 triggered SPM biosynthesis, we measured RvD1 in cell-free supernatants collected from CF MΦ stimulated with S1 protein. As shown (Fig. 2) after a 3 h treatment, RvD1 concentrations significantly increased compared to baseline in untreated samples, signifying activation of pro-resolutive SPM biosynthesis by leukocytes in response to SARS-CoV-2.

**Figure 2.**
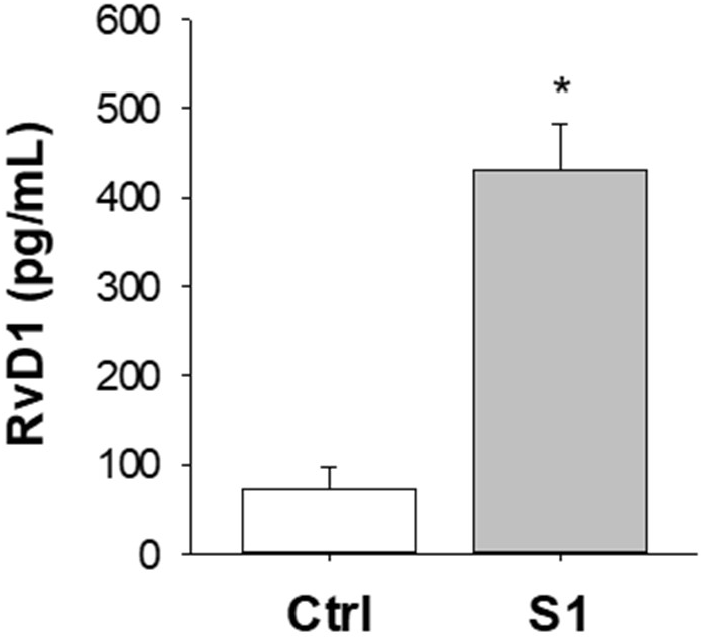
SARS-CoV-2 proteins stimulate RvD1 biosynthesis. RvD1 concentrations in MΦ cell supernatants following stimulation (3 h) with CoV-2 proteins (10 µg/mL). RvD1 was measured using a validated EIA procedure^14^. Results are mean ± SE from experiments with cells from 4 different donors. *, P < 0.05 vs untreated cells used as a control (Ctrl)

### SPM reduce CF MΦ inflammatory responses to SARS-CoV-2

Next we tested SPM bioactions on inflammatory responses triggered by SARS-CoV-2 proteins in CF MΦ. Since IL-8, which is crucial for neutrophil recruitment in acute inflammatory responses to infection, was increased in response to CoV-2 and SPM regulate neutrophil infiltration in inflamed tissue, we treated CF MΦ with S1, S2, and N proteins in combination with RvD1, RvD2, or LXA^4^ and determined if SPM treatment decreased IL-8 release by these cells. RvD1, RvD2, and LXA^4^ treatment resulted in a significant reduction in IL-8 protein in supernatants from CF MΦ treated with S1 or S2 and EtOH as a vehicle control, whereas only RvD1 and RvD2 decreased IL-8 in N-treated cells (Fig. 3). Since S1 protein plays important roles in SARS-CoV-2 first recognition by MΦ and cell infection, we focused on this viral protein and established effects of RvD1 and RvD2 on MΦ responses. RvD1 and RvD2 significantly reduced MCP-1 and MIP-1β that were increased by the S1 protein in CF MΦ and enhanced IL1RA thus countering the inhibitory effect of S1 on this cytokine. In addition, RvD1, but not RvD2, significantly dampened MIP-1α release and both gave a trend downward reduction in GM-CSF release (Fig. 4A). In addition, RvD1 and RvD2 significantly lowered TNF-α mRNA levels in S1-treated CF MΦ (Fig. 4B).

**Figure 3.**
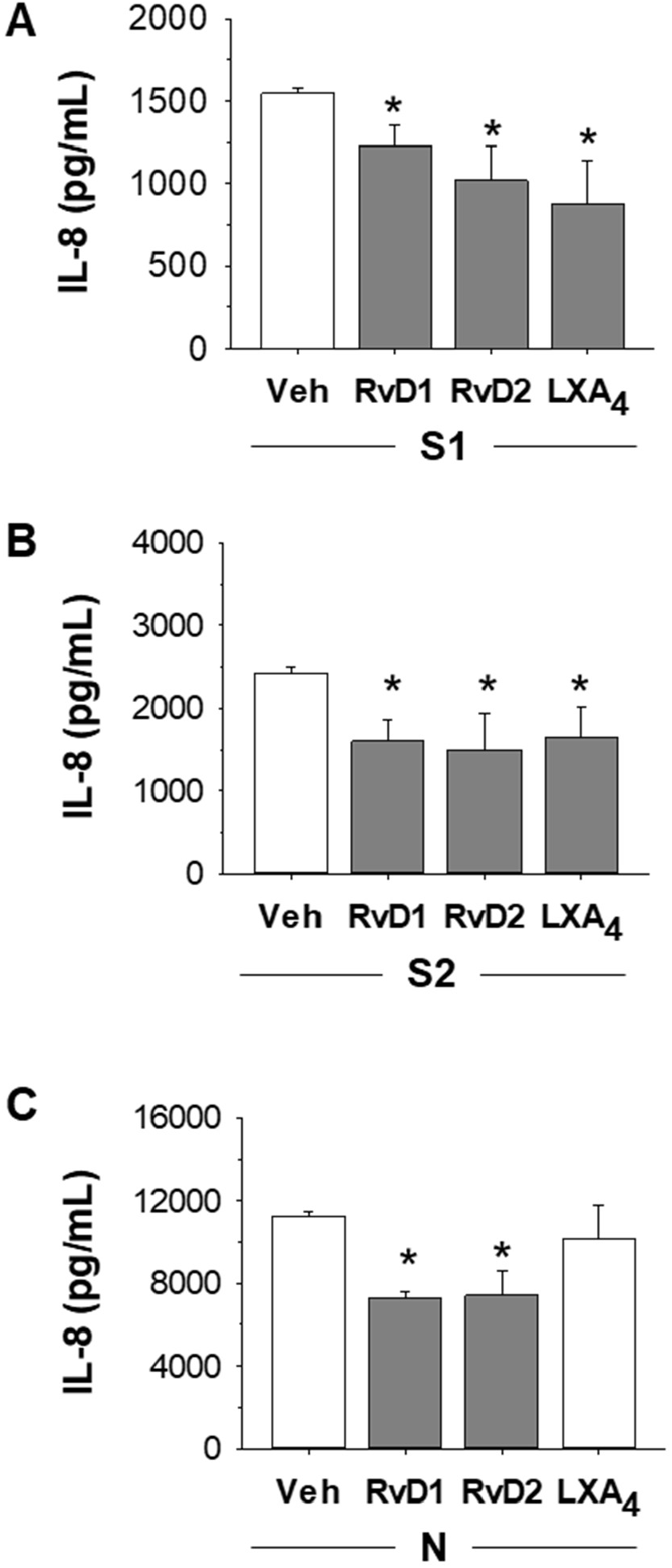
SPM actions on IL-8 release by CF MΦ in response to CoV-2. Monocyte-derived MΦ collected from volunteers with CF were treated with RvD1, RvD2, or LXA^4^ (10 nM) or EtOH (0.01 % vol/vol in PBS) as a vehicle (Veh) control and S1, S2, or N proteins (10 µg/mL, 3h). Cell free supernatants were harvested and used for measuring IL-8 release using a commercial ELISA. Results are mean ± SE from experiments with cells from 4 different donors. *, P < 0.05; **, P < 0.01; ***, P < 0.001 *vs* cells treated with CoV-2 proteins plus Veh.

**Figure 4.**
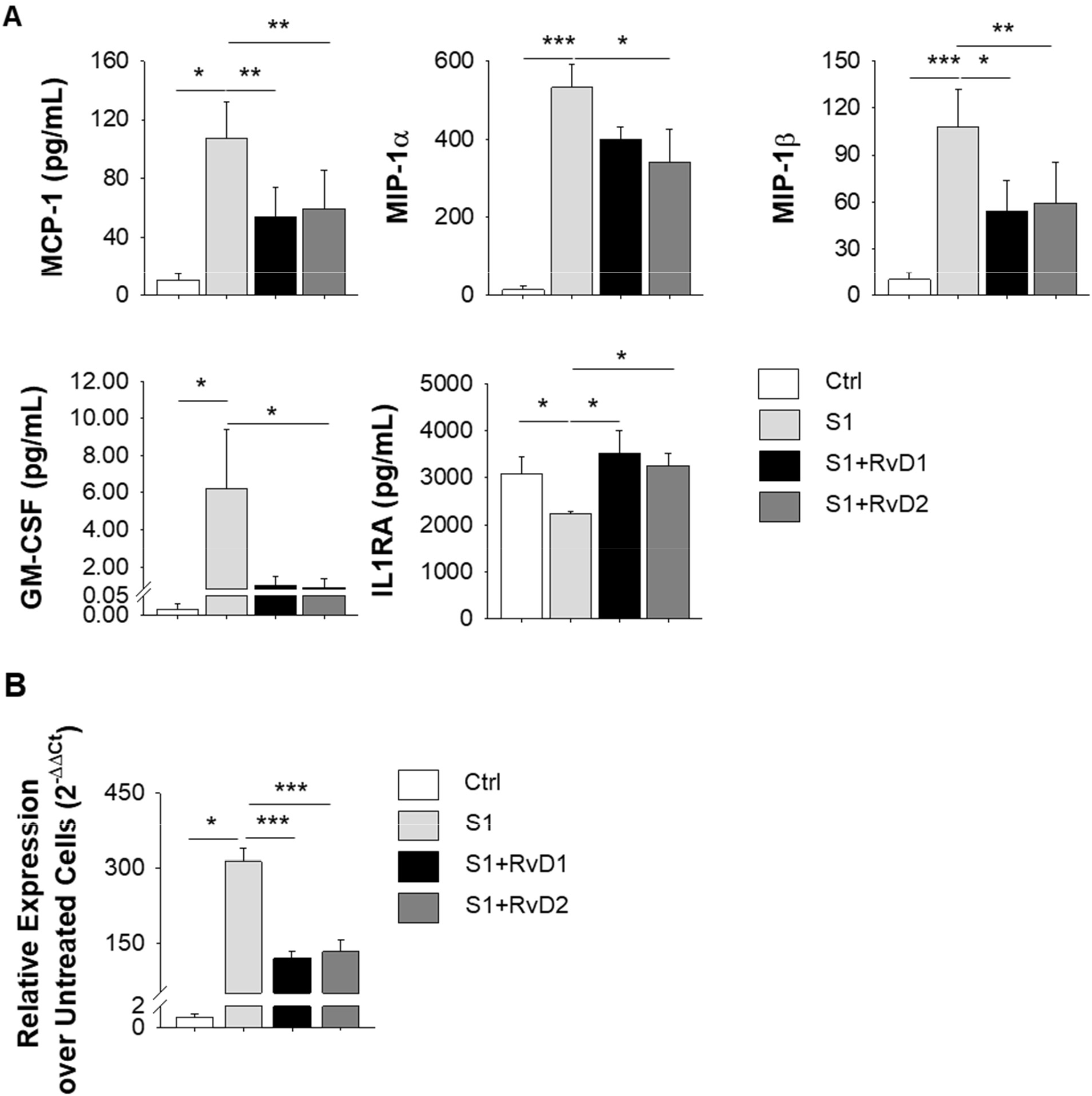
SPM actions on CoV-2 responses by CF MΦ. **(A)** Inflammatory mediators were determined in supernatants from monocyte-derived MΦ from volunteers with CF treated (3 h, 37 °C) with RvD1, RvD2 (10 nM), or Veh plus S1 (10 µg/mL) as described above. **(B)** Expression levels of TNF-α were determined with real time PCR analysis. Results are mean ± SE from experiments with cells from 4 different donors. *, P < 0.05; **, P < 0.01; ***, P < 0.001 *vs* cells treated with CoV-2 proteins and Veh.

Molecular mechanisms of action of SPM encompass regulation microRNAs that are important controller of inflammation. Therefore, given the previous findings we analyzed whether RvD1 and RvD2 modified miRNA in CF MΦ in response to SARS-CoV-2 S1 protein. Real time PCR analysis revealed that S1 protein significantly increased miR-197 while decreasing miR-16 and 29a (Fig. 5A). RvD1, but not RvD2, restored miR-16 and significantly reduced miR-197. Vice versa, RvD2 significantly enhanced miR-29a and miR-125a (Fig. 5B), demonstrating that each SPM has distinct regulatory activity on miRNA in MΦ in response to S1 SARS-CoV-2 protein.

**Figure 5.**
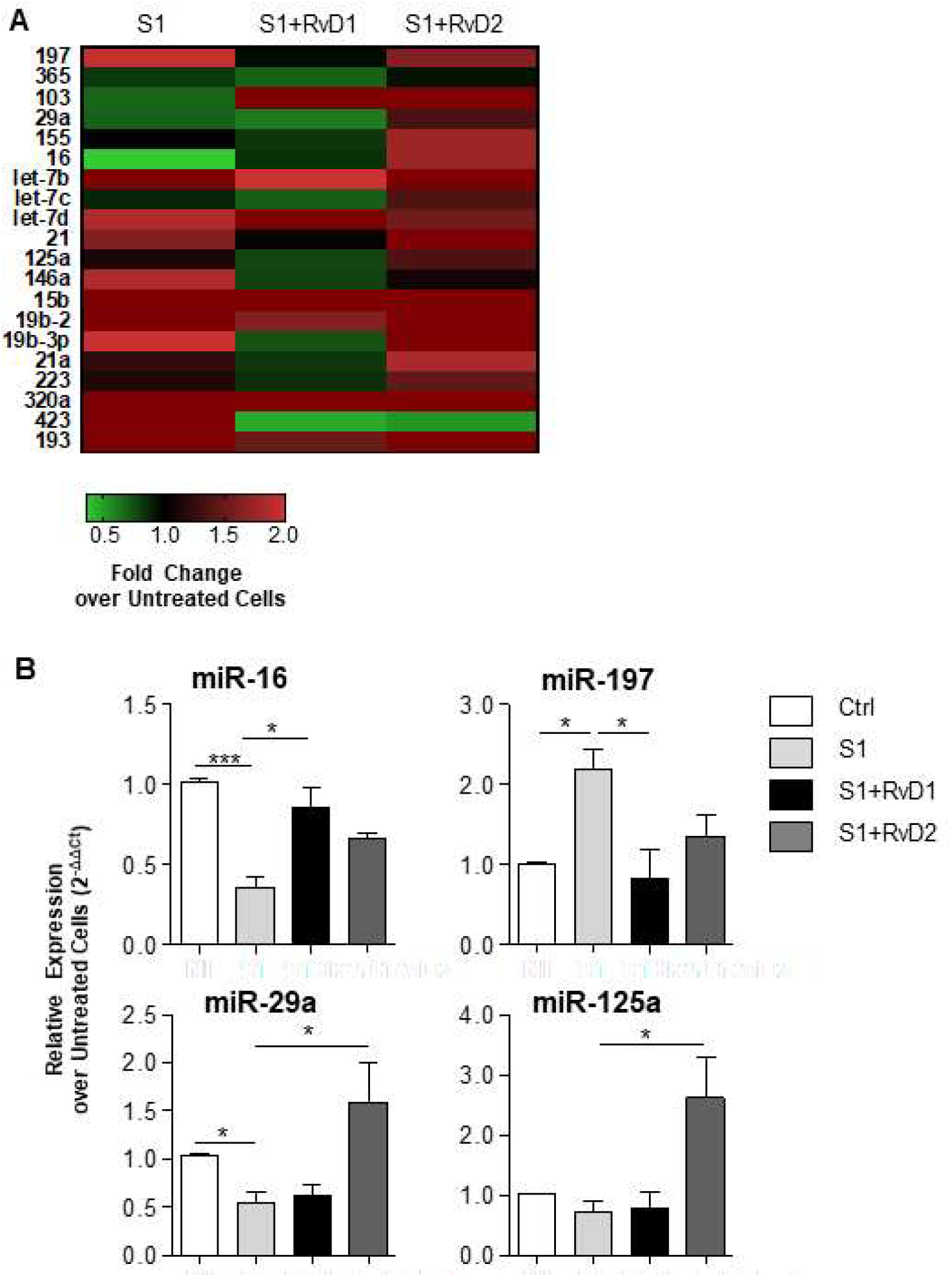
RvD1 and RvD2 regulates miRNAs expression in CF MΦ during CoV-2 response. Peripheral blood monocytes from volunteers with CF were isolated and treated with RvD1, RvD2 (10 nM), or vehicle as described above. S1 protein (10 µg/mL) was added (3 h). miRNA relative expression was determined and calculated using SNORD68 and SNORD95 as housekeeping miRNAs for loading control. The heatmap reports the average of fold change expression relative to untreated CF MΦ used as a control. Green squares indicate down-regulation, red squares indicate upregulation, black square no differences as compared to untreated CF MΦ. The histogram reports microRNAs relative expression levels as fold change over untreated CF MΦ. Results are mean ± SE from experiments with cells from 4 different donors. *, P < 0.05; ***, P < 0.001 b.

To determine the time-course of S1-induced responses and effects of RvD1 and RvD2, CF MΦ were treated with the S1 protein followed by RvD1, RvD2, or EtOH as a vehicle control after 3 h and concentrations of cytokines and chemokines were determined 24 h later. As shown, IL-8, TNF-α, and MIP-1α accumulated in MΦ supernatants and significantly increased from 3 to 24 h following S1 addition in a time-dependent manner, whereas IL-6, IL-1β, MIP-1β, and INF-α and γ did not change in S1-treated cells compared to unstimulated cells used as a control. RvD1 and RvD2 significantly diminished IL-8 (in keeping with results obtained with 3 h-treated MΦ) as well as TNF-α and MIP-1α (Fig. 6). Thus, IL-8 and MIP-1α, which are key leukocyte chemotactic signals, are rapidly and steadily induced in MΦ by S1 while TNF-α is activated as a second wave and protein levels of IL-6 and other cytokines or IFN involved in cytokine storm do not increase within 24 h from S1 stimulation despite a marked mRNA upregulation. These results also indicate that RvD1 and RvD2 reduce inflammatory responses and signaling to SARS-CoV-2 in CF MΦ.

## Discussion

Here, we report the characterization of cellular and molecular responses induced by SARS-CoV-2 in MΦ from individuals with CF and provide the first evidence for SPM biosynthesis and regulatory activities.

SARS-CoV-2 is a pleiotropic, highly contagious virus that can lead to an uncontrolled inflammatory syndrome in many organs especially in individuals with pre-existing chronic diseases like CF. MΦ are sentinel of host defense and key players in inflammation, resolution, and innate and adaptive immunity and have important role in CF. Therefore, here we chose MΦ as a cell model for determining inflammatory actions of SARS-CoV-2 in CF. While infection with whole virions is important to assess overall viral replication and host responses, the use of single proteins represents a useful strategy for discriminating the contribution of each viral component to inflammation in target cells. The host innate immune system recognizes SARS-CoV-2 proteins through pathogen recognition receptors (PRR) including toll-like receptor (TLR) 4 and NOD-like receptors (NLR) that form the inflammasome machinery. Recognition and activation of TLR4 culminate with the activation of the NF-κB and interferon-regulatory factors (IRF) leading to the subsequent production of NF-κB-dependent cytokines and chemokines and IFN, respectively. Here we found a rapid increase in the release of chemokines (IL-8 being predominant) but not of cytokines (IL-1β, IL-6) or INF-α, β, and γ in response to short (3 h) stimulation of CF MΦ with S and N proteins and, in response to prolonged (24 h) S1 treatment, TNF-α was the only cytokine increased (Fig. 1 and 6). These results demonstrate a temporal dissociation in MΦ responses to SARS-CoV2 where there is an early induction of chemokines that stimulate recruitment neutrophils and other leukocytes and a delayed secretion of TNF-α that can lead to MΦ self-activation. Our findings on IL-8 also prove that its rapid induction does not occur at transcription level. These results also indicate that some NF-κB-responsive gene (e.g., IL-6), along with the IRF/IFN and inflammasome pathway, are not efficiently activated by S1. This was particularly intriguing for IL-6, since its protein levels were not increased even following a 24 h stimulation and despite a ∼ 200 fold upregulation of mRNA. In in vitro studies, Cheung and coworkers found that SARS-CoV did not induce IFN-α/β secretion in human MΦ in contrast with H1N1 influenza virus and suggested that this deficiency in anti-viral response could explain some of the clinical manifestations of SARS-CoV infection^23^. It is possible that multiple signals (e.g., co-stimulation of cell-membrane and phagosome-associated PRR) that occur in vivo during SARS-CoV-2 infection are required to induce secretion of IL-6 and IFN.

SPM are rapidly biosynthesized from polyunsaturated fatty acids in sterile and infectious inflammation. Here were report that SARS-CoV-2 proteins trigger the biosynthesis of RvD1 in peripheral blood and isolated MΦ from volunteers with CF (Fig. 2), which is paradigmatic of activation SPM biosynthetic pathways. SPM precursors, including 17-HDHA (the precursors RvD1 and RvD2) have been identified in lipidomic analyses from mouse airway lavage fluids and human nasal washes during H1N1 influenza ^24^. Moreover, 17-HDHA and its derivative protectin D1 proved to have anti-viral and vaccine-adjuvant activities^13,17^. In this work, we found that DHA-derived RvD1 and RvD2 and the AA-derived LXA^4^ have counter-regulatory actions on SARS-CoV-2-induced inflammatory responses in CF MΦ, including reduction of select inflammatory chemokines and cytokines (Fig. 3, 4, 6). SPM are potent regulator of neutrophil chemotaxis and recruitment in inflamed tissues. For instance, we recently demonstrated that RvD1 reduces IL-8 in MΦ from volunteers with CF and KC (the murine IL-8 homologue) CF mice infected with *P. aeruginosa* ^14^ whereas RvD2 curbs chemokine storm in septic mice (7). RvD1 also increased expression level of miR-16 and reduced miR-197 that were modified by S1 in CF MΦ, whereas in RvD2-treated cells we observed a significant increase in miR-29a and miR-125a (Fig. 5). Noteworthy the identified microRNAs regulated by RvD1 and D2 have roles in the inflammatory response and inflammatory diseases. miR-16 and miR-125a are negative regulators of NF-κB activation, reduce cytokine/chemokine production in MΦ, limit leukocyte infiltration in lungs in vivo^25–27^; miR-29a acts as a tumor suppressor in acute myeloid leukemia (30) and its overexpression results in a significant reduction in TNF-α release by stimulated human MΦ^29^; miR-197 amplifies cytokine signaling and it has been proposed as a therapeutic target for reducing tumor-associated inflammation^30^. Regulation of NF-κB and NF-κB-associated microRNAs is a central mechanism of action of SPM in resolution. RvD1 blocks NF-κB nuclear translocation^31,32^, upregulates miR-146b in human MΦ that results in downregulation of inflammatory cytokines and chemokines^32^. In vivo, RvD1 dampens NF-κB activation and downstream genes in lungs and kidneys from sickle cell mice undergoing hypoxia-reoxygenation injury^33^, enhances miR-219 in mouse peritoneal leukocytes leading to increase in IL-10 (37), and increases miR-21 and miR-155 that interact with the TLR-NF-κB axis in lung MΦ from mice bearing chronic *P. aeruginosa* infection (22). RvD2 induces miR-146a in human monocytes stimulated with LPS attenuating TLR-mediated inflammation^34^. Of interest, reduced expression of the RvD1-regulated miR-219 has been associated with impaired resolution of inflammation (1), corroborating the biological importance of SPM-regulated microRNAs in the regulation of inflammation. Hence, results shown here give the first evidence for roles of miRNAs in SPM regulation of inflammatory responses to viral stimuli.

In summary, here we show that SARS-CoV-2 triggers inflammatory responses in MΦ from patients with CF that encompass rapid secretion of a broad number of leukocyte chemokines and delayed induction of select cytokines that can amplify MΦ hyper-inflammation. Moreover, we demonstrate that SARS-CoV-2 initiates the biosynthesis of SPM including RvD1 that, along with and RvD2, control MΦ responses related to NF-κB activation and cytokine/chemokine production by targeting miR-16, miR-197, miR-29a, and miR-125a.

